# A cross-sectional study to assess the magnitude of Hypertension and Type 2 Diabetes Mellitus in Hatcliffe, Harare

**DOI:** 10.1101/535450

**Authors:** Lonestar Lazarus Gonde, Moses John Chimbari, Tawanda Manyangadze

**Author notes:** **Corresponding Author: Lonestar Lazarus Gonde**, University of KwaZulu Natal, College of Health Sciences School of Nursing and Public Health, 238 Mazisi Kunene Rd, Glenwood, Durban, 4041, South Africa.

## Abstract

**Background:** Hypertension (HTN) and type 2 diabetes mellitus (T2DM) are reported to be on the increase in developing countries. In this study we investigated the epidemiology of the prevalence of HTN and T2DM and its correlates in a high density area. We carried out this study to assess the magnitude of the prevalence of HTN and T2DM so that we can query the drivers that are causing an increase conditions in these conditions.

**Methods:** We conducted a cross-sectional study in Hatcliffe, a high density area (HDA) in Harare. We interviewed, bled, took anthropometric and measured blood pressure of 381 participants. We investigated HbA1c, blood pressure, BMI and prevalence of prehypertension, prediabetes, co-existence of HTN and T2DM. A geospatial analysis was carried out to ascertain distribution patterns of HTN and T2DM in Hatcliffe.

**Results:** The prevalence for prehypertension and prediabetes is higher than it is for full blown conditions of the HTN and T2DM. The prevalence of prehypertension was 35.4% and prediabetes was 29%. The prevalence of HTN in this study was 14.4% whilst that of T2DM was 3.93%. Out of the 55 participants that had developed HTN five had developed T2DM. There was no significant difference in the HTN and T2DM clusters.

**Conclusions:** The prevalence for prehypertension and prediabetes is higher than it is for full blown conditions of the HTN and T2DM. This indicates the importance of having a strategy for reducing the number of prediabetes and prehypertensive cases so that cases of full blown T2DM and HTN remain low.

## Introduction

Hypertension (HTN) and type 2 diabetes (T2DM) are important non-communicable diseases (NCD) which are on the increase globally but more so in low to medium countries (LMIC) particularly in sub-Saharan Africa (SSA) (1–3). HTN and T2DM have been referred to as a “slow motion public health disaster” mainly because very little attention is paid towards them the damage they would further create in the future would be immense(4). Until recently HTN and T2DM were seen as diseases that only affect the rich (5).

Both HTN and T2DM have a characteristics of being asymptomatic which results in low awareness leading to many cases remaining undiagnosed and contributing to the high mortality in SSA (6–8). The prevalence of the ‘toxic combination’ of HTN and T2DM was reported to be equal to T2DM alone in Cameroon (9).HTN and T2DM are important risk factors for cardiovascular diseases which overlap and have a common pathophysiology (9–11). Hypertension is common in individuals that have already developed T2DM(12). Projections indicate that by 2035, SSA will have 41 million people with T2DM(13). Other projections indicate that by 2025, 125million people in SSA will have HTN (14). In a study carried out in Ghana it was found that the weighted prevalence of prehypertension was 30% (24). Prediabetes in SSA is projected to increase by 75.8% to 47.3 million in 2030 from the 26.9 million that were reported in 2010 (26).

There is paucity of information on HTN and T2DM in some SSA countries and that makes it difficult to develop appropriate control strategies that would reduce the prevalence (8, 15). The coexistence of HTN and T2DM increase the cost of health for individuals and at a national level(16). Information on the prevalence of prehypertension and prediabetes is also scarce in SSA (8, 17).

The majority of people that succumb to HTN and T2DM in SSA reside in the urban areas (18). Furthermore urbanization, sedentary lifestyle and nutritional transition contribute to the increase in the prevalence of HTN and T2DM (5, 6, 19).

In this study we investigated the epidemiology of the prevalence of HTN and T2DM and its correlates in a high-density area (HDA) in Harare, Zimbabwe. The results of this study will bring better understanding of the magnitude of HTN and T2DM which will better inform relevant stakeholders such as the City Council and the Ministry of Health and Child Care to improve primary healthcare in the HDA.

## Research methods and Design

### Study design and population

We carried out a cross-sectional study in Hatcliffe, a high-density area HDA situated 17° 41’ 18” South, 31° 6’ 35” East. Hatcliffe is the second fastest growing HDA in Harare with an estimated population of 5000 people. A HDA in Zimbabwe is defined as an areas with residential stands that have an average of 300 square meters. Hatcliffe is subdivided into different sections with different names. Some of these sections in Hatcliffe are still developing. Two of the oldest sections of Hatcliffe, namely Cooperative and Old Hatcliffe are built up and have little room for expansion.

### Inclusion and exclusion criteria

We interviewed (face to face) consenting participants who had resided in Hatcliffe for more than six months. We interviewed a selected male and a female aged 18 to 75 from every fourth house from the ‘Old Hatcliffe ‘ and ‘Cooperative’ areas. We excluded vulnerable people such as pregnant women and mental patients. Bed-ridden patients and those with confirmed renal failure were also excluded from the study. Participants agreed to take part in the study by way of appending their signature on the informed consent form after explaining to them details of the study and the consequences of their participation. A geospatial analysis was carried out to ascertain distribution patterns of HTN and T2DM in Hatcliffe.

### Sample size

Since the prevalence of the HTN and T2DM was not known we calculated the sample using the formula N = Z2 P (1-P)/e2 as informed by Stepwise NCD studies. N is the sample size, Z is the level of confidence, P is the estimated proportion of an attribute that is present in the population. P was estimated at 0.50; Z = 1.96 and e = 0.05, thus the estimated sample size was N = 1.962 × 0.5(1−0.5)/0.052 = 384 and a precision level of 5% at 95%confidence was used.

### Data collection

Data was collected by the project principal investigator and four professional community mobilizers from Hatcliffe between 21 September 2018 and 4 October 2018. The community mobilizers had received training in KoBo Toolbox. A questionnaire adapted from the World Health Organization NCD STEPS survey was used for the face to face interviews conducted at household level. The questionnaire was loaded into a hand held Android mobile device installed with Kobo Toolbox.

Participants were asked to present themselves at the clinic so that their blood pressure could be measured. Ten milliliters of blood where taken from each participant for the purposes of investigating total HbA1c, creatinine and total cholesterol. Creatinine levels can determine if a participant has renal disease or not.

Anthropometric measurements recorded for each participant. Participants whose body mass index (BMI) was between ≥25.0 and 29.9 were defined as overweight whilst a BMI ≥30.0 indicated being obese. A BMI between 18.5 and 24.9 kg/m2 was defined as normal. Participants with a BMI of less than or equal to 18.5kg/m2 were categorized as underweight. A calibrated Salter weighing scale was used to measure weight and a stadiometer was used to measuring height. Three blood pressure readings were taken at 5 minutes intervals using a digital Health Easy blood pressure machine. For the purposes of this study we used JNC standards which define hypertension as systolic BP ≥ 140 mmHg and/or diastolic BP ≥ 90 mmHg. We further designated participants into prehypertension, stage one and two hypertension groups as shown in Table 1. Being pre-hypertensive alerts clinicians if a patient is at risk of hypertension or not. Patients categorized as stage one or two of hypertension need to be put on treatment for hypertension.

**Table 1:** Categories of blood pressure according to the Joint National Committee

| **Blood pressure Classification** | **Systolic Blood Pressure** | **Diastolic Blood Pressure** |
| --- | --- | --- |
| **Normal** | < 120 | and < 80 |
| **Prehypertension** | 120-139 | or 80-89 |
| **Stage 1 Hypertension** | 140-159 | or 90-99 |
| **Stage 2 Hypertension** | $\geq 160$ | or $\geq 100$ |

**Table 2:** Frequency distribution of sociodemographic factors of study participants

| Variables | Total, n(%) | Female, n(%) | Male, n(%) | P -value |
| --- | --- | --- | --- | --- |
| **Age** |  |  |  |  |
| 18-24 | 101(26.5) | 39(19.9) | 62(33.5) | 0.309 |
| 25-34 | 94(24.7) | 52(26.5) | 42(22.7) |  |
| 35-44 | 81(21.3) | 45(23.0) | 36(19.5) |  |
| 45-54 | 46(12.1) | 26(13.3) | 20(10.8) |  |
| 55+ | 59(15.5) | 34(17.4) | 25(13.5) |  |
| **Level of Education** |  |  |  |  |
| Never been to school | 21(5.5) | 17(8.7) | 4(2.2) | 0.051 |
| Received some primary education | 3(0.8) | 2(1.0) | 1(0.5) |  |
| Finished primary education | 35(9.2) | 22(11.2) | 13(7.0) |  |
| Finished O level education | 229(60.1) | 110(56.1) | 119(64.3) |  |
| Did not finish O level education | 57(15.0) | 35(17.9) | 22(11.9) |  |
| Finished A level education | 20(5.3) | 6(3.1) | 14(7.6) |  |
| Did not finish A level education | 1(0.3) | 0 | 1(0.5) |  |
| Finished tertiary education | 15(3.9) | 4(2.0) | 11(6.0) |  |
| **Marital status** |  |  |  |  |
| Single | 99(26.0) | 20(10.5) | 79(41.6) | <0.001 |
| Married | 240(63.0) | 142(74.4) | 98(51.6) |  |
| Divorced | 15(3.9) | 6(3.1) | 9(4.7) |  |
| Widowed | 23(6.0) | 20(10.5) | 3(1.6) |  |
| Cohabiting | 4(1.0) | 3(1.6) | 1(0.5) |  |
| **Income, median(IQR)** | 0(0-150) | 0(0-70) | 50(0-200) | <0.001 |
| **Income groups** |  |  |  |  |
| 0-50 | 232(60.9) | 134(70.2) | 98(51.6) | <0.001 |
| 51-100 | 45(11.8) | 28(14.7) | 17(9.0) |  |
| 101-250 | 60(15.8) | 18(9.4) | 42(22.1) |  |
| 251-500 | 41(10.8) | 10(5.2) | 31(16.3) |  |
| 500+ | 3(0.8) | 1(0.5) | 2(1.1) |  |
| **Employment status** |  |  |  |  |
| Student | 6(1.6) | 0 | 6(3.2) | <0.001 |
| Not employed | 206(54.1) | 123(64.4) | 83(43.7) |  |
| Self employed | 53(13.9) | 15(7.9) | 38(20.0) |  |
| Formally employed | 95(24.9) | 45(23.6) | 50(26.3) |  |
| Retired | 21(5.5) | 8(4.2) | 13(6.8) |  |

### Biochemical

#### HbA1c

In this study T2DM was defined as HbA1c of 6.5% or above. Participants with an HbA1C between 5.7% and 6.4% were classified as prediabetes. A reading of 5.6% meant that the participant was not diabetic.

#### Specimen collection and preparation for HbA1

The pretreatment procedure for whole blood specimens involved initial centrifugation of specimens at 2000rpm per minute then aspiration of 25 μl of corpuscle from the blood corpuscle deposition into a microfuge tube. 500 μl of pre-treatment solution was added to the microfuge tube and closed before lysing the blood by vigorously shaking. The hemolysate was used as the working sample after sitting for 5 minutes.

#### Total Cholesterol

We classified a total cholesterol level between 5.17 mmol/l and 6.18 mmol/l as borderline high whilst ≥ 6.21 mmol/l was considered high. A cholesterol level of less than 5.17 was considered to be normal.

Serum samples were used for the total cholesterol assay on the Mindray BS 200E chemistry analyser. Specimens were centrifuged at 3000xg for 15 minutes and stored at −20 degrees Celsius.

### Data analysis

Data was collected using the KoBo Toolbox database before being exported to Excel for management. STATA 15 was used for all analyses in this study. Categorical variables were presented as proportions compared by gender. Level of significance in all analysis was set at P < 0.05 at 95% confidence interval. Bivariate analysis was used to consider significant variables (< 0.05) and a multivariate logistic regression model based on significant bivariate variables was performed to determine adjusted risk factors for HTN and T2DM. Hypertension and T2DM spatial clusters were determined by way of Kulldorff’s scan statistic in SaTScan™ (20, 21). The Kulldorff’s scan statistic tests the randomness of the events such as diseases cases in space and is able to specify the approximate location of significant and not significant clusters (22). The Bernoulli method in SaTScan™ uses the “0”and “1” event data to detect prevalence clusters. We used this approach in our study as used by Manyangadze et al (23). The cases of hypertension and/or diabetes were represented by “1” and the negative cases were represented by “0”.). The cases and negatives were used to determine if there was clustering of the case location distribution relative to the negatives location distribution (24) SaTScan™ uses a moving window of varying diameter to detect and evaluate clusters. In this study we calculated the number of observed and expected observations, percentage cases in area and relative risk. The SaTScan Bernoulli model uses a likelihood ratio testthe probability of a group of cases within a potential cluster defined by a circle. Relative risk is the estimated risk within the cluster divided by the estimated risk outside the cluster. In this study we report all the non-overlapping. These high-risk clusters were based on maximum radius of one kilometer (considering the size of the study area), 50% of the total number of subjects. We did not adjust for more likely clusters. The 999 Monte-Carlo replications was conducted to evaluate statistical significance of the clusters (22).

### Ethical approval

Ethics approval to conduct the study was obtained from Medical Research council of Zimbabwe (A/2281) and Biomedical Research Ethics Committee of the University of KwaZulu-Natal (BE628/17). Gatekeepers’ permission was sought from Harare City Council Health Department as well as Ministry of Health and Child Care, Zimbabwe.

## Results

The socio-demographic characteristics of the participants are shown in Figure 2. Females constituted 51.4 %(n=196) of the study participants whilst 48.6% (n=185) were males. The majority (84.5 %) of the participants had reached a level of education of ‘O’ level or higher (p-=0.051). Two hundred and forty of the participants were married (p<0.001). The majority of the participants earned less than $50 per month (p< 0.001). More than half (54.1%) of the participants were not employed (p<0.001). Twenty nine point nine percent (29.9%) of the participants said they consumed alcohol and of these seventeen (17) were women (p< 0.001).

The overall prevalence of HTN among the study participants of Hatcliffe was 14.4% (p=0.099) as shown in Table 3. Out of the 55 confirmed cases of HTN, 32 females and 23 males had full blown HTN. On the other hand the prevalence of T2DM of the study participants was 3.93% (p=0.095). Ten females and 5 males had developed T2DM.

**Table 3.** Prevalence of Hypertension and Type 2 Diabetes Mellitus by sex

| Condition | Total |  | Female |  | Male |  | p-value |
| --- | --- | --- | --- | --- | --- | --- | --- |
|  | n | p[CI] | n | p[CI] | n | p[CI] |  |
| Hypertension | 55 | 0.144[0.11-0.18] | 32 | 0.17[0.12-0.23] | 23 | 0.12[0.08-0.18] | 0.099 |
| T2DM | 15 | 0.039[0.02-0.06] | 10 | 0.05[0.03-0.09] | 5 | 0.03[0.01-0.06] | 0.095 |

Out of the 381 participants, 60 females and 75 males were at the pre-hypertensive stage as shown in Table 4. Twenty nine percent of the participants were at a pre-diabetic stage. Eighteen (18) males and 11 females were reported to be pre-diabetic.

**Table 4:** Frequency distribution of anthropometric and clinical characteristics for study participants

| Variables | Total, n(%) | Female, n(%) | Male, n(%) | P -value |
| --- | --- | --- | --- | --- |
| Blood pressure |  |  |  |  |
| Normal | 191(50.1) | 99(51.8) | 92(48.4) | 0.253 |
| Prehypertensive | 135(35.4) | 60(31.4) | 75(39.5) | 0.05 |
| Stage 1 | 40(10.5) | 23(12.0) | 17(9.0) | 0.163 |
| Stage 2 | 15(3.9) | 9(4.7) | 6(3.2) | 0.218 |
| HbA1c |  |  |  |  |
| Negative | 337(88.5) | 170(89.0) | 167(87.9) | 0.183 |
| Prediabetes | 29(7.6) | 11(5.8) | 18(9.5) |  |
| Diabetes | 15(3.9) | 10(5.2) | 5(2.6) |  |
| BMI |  |  |  |  |
| Normal | 219(57.6) | 101(53.2) | 118(62.1) | 0.039 |
| Overweight | 77(20.3) | 48(25.3) | 29(15.3) |  |
| Obese | 33(8.7) | 23(12.1) | 10(5.3) |  |
| Underweight | 51(13.4) | 18(9.5) | 33(17.4) |  |
| Cholesterol levels |  |  |  |  |
| Normal | 316(83.6) | 155(81.6) | 161(85.6) | 0.143 |
| Borderline | 45(11.9) | 27(14.2) | 18(9.6) |  |
| High | 17(4.5) | 8(4.2) | 9(4.8) |  |

Hundred and ten (110) participants were obese with the majority of them being women. Borderline cholesterol levels were detected in 45 participants of which 27 were women.

Five (9.1%) out of 55 hypertensive participants were diabetic (Table 5). Diabetic individuals were three point five times more likely to develop HTN and individuals with high cholesterol levels were 2.06 more times likely to develop HTN.

**Table 5:**
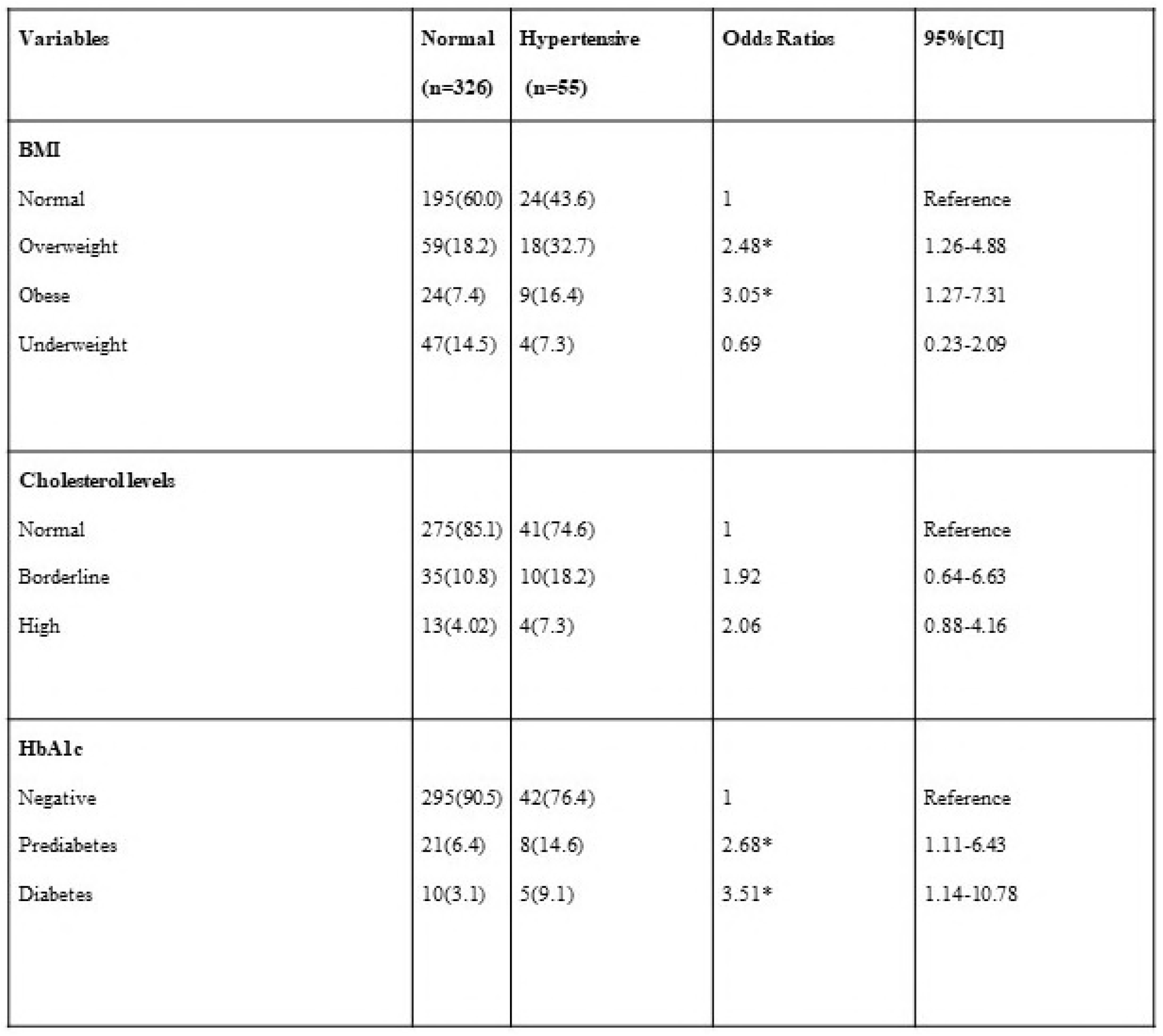
Bivariate analysis of medical factors and hypertension among study participants.

**\* Significant independent variable**

Based on the cluster analysis Figure 1 shows that the distribution of HTN was haphazard.

**Figure 1:**
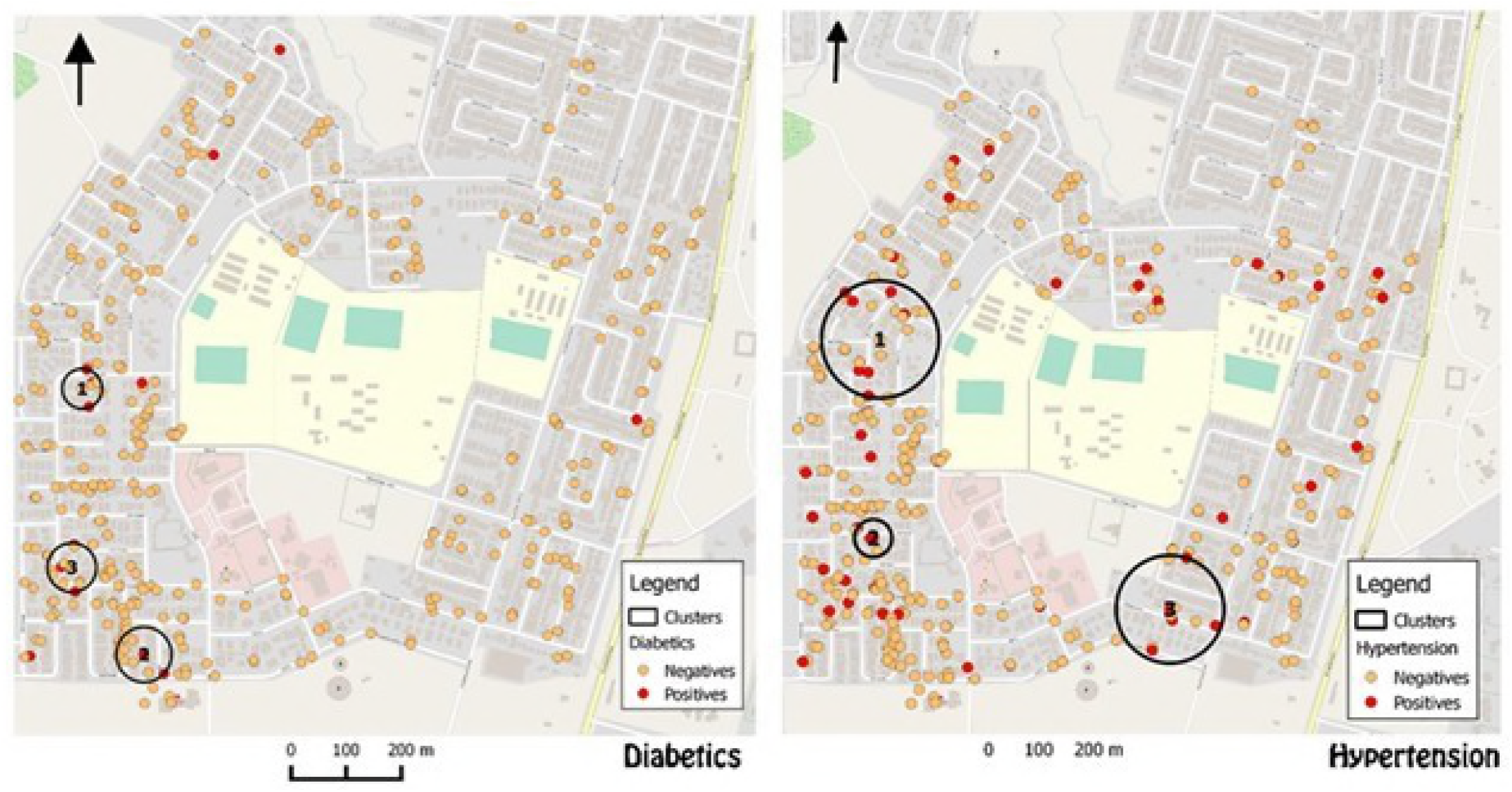
Map of Hatcliffe showing positions of clusters (circled) of type 2 diabetes and hypertension.

Furthermore there were no differences between the clusters and areas outside clusters (Table 6).

Table 6: Properties of the Hypertension and Type 2 Diabetes clusters in Hatcliffe suburb in Harare, Zimbabwe

## Discussion

The prevalence of HTN and T2DM in Hatcliffe HDA was lower than the national estimations. Mutowo suggests that the estimated national prevalence for Zimbabwe for T2DM is 5.9% whilst the pooled prevalence for HTN is 30% (21, 22). However Mutowo’s study used pooled secondary data collected using different methodologies with different quality checks. Our study used primary data which allowed us to do quality checks. Thus, we believe that our results are accurate. It is also possible that the prevalence of T2DM detected in our study is more accurate because the HbA1c test is much more sensitive than the fasting plasma glucose employed in many primary healthcare settings in Zimbabwe (25).

The prehypertension prevalence of 35.4% exhibited in the study is higher than overall age-adjusted prevalence of 25.9 % found in a four sub-Saharan country study (8). The pre-hypertension prevalence in this study may be used to forecast the future actual prevalence. Should all the pre-hypertensive cases identified in this study eventually convert to actual confirmed cases there would be a doubling of the current prevalence. In this study, the percentages of participants that had already developed HTN and those that were prehypertensive almost matched those without HTN (n=49.9 vs n=50.1). The implications are that if all those prehypertensive eventually convert to full blown hypertension there could be serious problems for Hatcliffe. In our study there were more pre-hypertensive males than females. Our findings are consistent with findings from a nationwide study in Nepal, carried out by Agho *et al* which reported the number of men with prehypertension to be higher than that of women (24).

To the best of our knowledge very few studies have determined the prevalence of prediabetes in Zimbabwe. In this study there were more pre-diabetic males than females. Knowing that more males than females are pre-diabetic in Hatcliffe could better prepare for programs to reduce or reverse the risk factors that are gender specific.

A few studies have been conducted in SSA to ascertain the level of coexistence of HTN and T2DM. Most studies based on known hospitalized diabetic patients reported the prevalence of the ‘toxic combination’ of HTN and T2DM to be equal to T2DM alone in Cameroon (9). Hospitalized patients are easier to reach compared to participants in the community. However, such patients do not truly represent what is in the population from which they come. More needs to be done to determine the coexistence of HTN and T2DM in a community setting. Our study is one of the few that have looked at co-existence of HTN and T2DM in SSA at a community household level.

We cannot conclusively postulate that there was clustering because clusters were not significant. Instead, we can confirm that the development of HTN or T2DM was haphazard, occurring randomly across the study area. Therefore, the probability of getting someone with the disease or condition is the same across the study area. Hence the treatment or awareness campaigns should be area wide rather than targeting specific areas.

## Conclusion

We conclude that that prevalence of pre-hypertension and pre-diabetes is higher than that of independent HTN and T2DM. We also established that prevalence of the co-existence of HTN and T2DM was very low. To avoid pre-diabetic and pre-hypertensive cases from progressing to full blown HTN and T2DM, a strategy for earlier identification of the pre-diabetic and pre-hypertensive cases should be implemented. More geospatial studies are needed to link the confirmed cases of HTN and T2DM with risk factors. Given that we only used one method for cluster analysis, it may be worthwhile to use complementary methods to determine clustering and detect hotspots.

## Acknowledgments

We would like to thank the statistician Tafadzwa Madanhire, the team of data collectors, PCD Wholesalers, Greenwood Wholesalers and Providence for their support. We would like to acknowledge financial support from UKZN Colleges of Health Scholarship.

## Competing interests

None

## Author Contributions

Wrote the first draft of the manuscript and collected the data: LLG. Contributed to the writing of the manuscript: LLG, MJC and TM. Data analysis, interpretation of the manuscript: MC, TM, LLG. Criteria for authorship read and met: LLG. Agree with manuscript results and conclusions: LLG, MC,TM.

